# A Database of Restriction Maps to Expand the Utility of Bacterial Artificial Chromosomes

**DOI:** 10.1101/2023.03.31.535162

**Authors:** Eamon Winden, Alejandro Vasquez-Echeverri, Susana Calle-Casteneda, Yumin Lian, Juan Pablo Hernández-Ortiz, David C. Schwartz

## Abstract

While Bacterial Artificial Chromosomes were once a key resource for the genomic community, they have been obviated, for sequencing purposes, by long-read technologies. Such libraries may now serve as a valuable resource for manipulating and assembling large genomic constructs. To enhance accessibility and comparison, we have developed a BAC restriction map database.

## Background

Bacterial Artificial Chromosomes (BACs) were a central resource for early sequencing and mapping efforts that enabled construction of the first reference genome for human and other higher eukaryotes [1, 2]. While BACs have been largely supplanted for creating reference genomes [3, 4], research directions in synthetic biology are now advancing towards large-scale genome alterations and assemblies. Consequently, construction technologies are emerging as an increasingly valuable resource for contemporary investigators [5]. Accordingly, BACs could be leveraged as expansive, well-characterized genomic elements to be used as building blocks. In this regard, there are currently limited resources available for fully engaging the many advantages offered by BACs. Consider that while there are fully sequenced BACs, including those in the “golden path” used to generate drafts of the human genome, the UCSC Genome Browser presents a collection of end-sequenced BACs [6] and NCBI’s cloneDB that has been discontinued [4]. Consequently, synthetic efforts would benefit from resources that readily functionalize BACs as physically manipulable molecules, or “parts,” for genome writing activities. One such fundamental resource is a database of restriction maps created from available human BAC clones.

### Data Description

BAC DNA molecules require special considerations for synthetic workflows due to their large size and for the mediation of cloning and sequencing errors—particularly when dealing with complex, repeat-ridden portions of the human genome [7, 8]. Here, a comprehensive set of restriction maps simplifies selection and validation of optimal BACs within libraries for such applications.

### Analyses

We created this resource using publicly available data from National Center for Biotechnology Information (NCBI)[9]. End-sequenced BACs with unique placement were mapped, as were insert-sequenced BACs. The resulting database of restriction maps includes most BACs that are publicly available for the human genome. As sequencing costs have reduced dramatically, the later addition of more libraries or species would be welcome. The diversity of BAC libraries, assembled from different donors and by different methods, enriches the diversity and utility of available sequences for such uses as genome writing applications. This resource offers restriction maps available across many BACs enabling systematic selection of clones and restriction fragments for a broad range of applications.

## Discussion

This database establishes a useful resource for directed manipulations of large insert clones (BACs) as a convenient way to explore sizable, discrete chunks that would readily allow the contextualization of genic regions within non-coding, genomic “dark matter” portions of the genome. With ready access to comprehensive restriction maps, searching for BACs with specific characteristics or developing workflows concerning linearization and vector removal are now enhanced and simplified. In Figure 1, you can see an output of a simple pipeline determining a clone and fingerprint map from a region of interest and then visually representing it. CRISPR-Cas systems have revolutionized this space and pairing this resource with one of the many guide-RNA libraries to find targets for manipulation with CRISPR tools further synergizes BAC advantages for genomic research.

**Figure 1:**
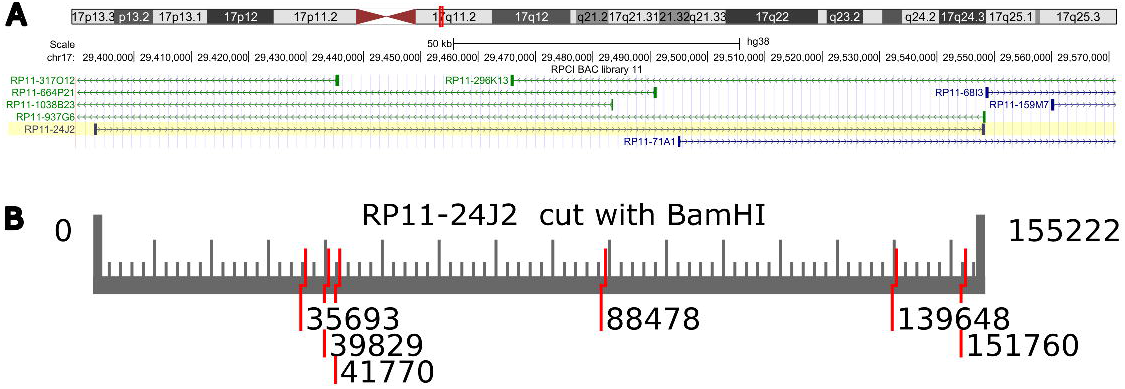
A) A screenshot of the UCSC genome browser (http://genome.ucsc.edu[6]) where the RP11-24J2 clone is placed in chromosome 17 of the human genome. B) An example of the insert of RP11-24J2 mapped by BamHI and drawn with a Python function, drawMap, which is included in the package, bacmapping, that was written for this database. The 5’ overhangs are shown to improve clarity.

### Potential Implications

BACs are a versatile resource for well-characterized genomic fragments that have been created for a broad range of organisms [10]. While these libraries have served well to establish reference genomes, they may now be refashioned as a source of large-scale building blocks for genome construction and manipulations where they will empower new ways to investigate chromosome-scale biology [11]. These studies require a resource of BAC components, ready to be forged into final products. Accordingly, databases like this one provides the basic information scientists need to start selecting BACs from the available libraries.

## Methods

Sequence and details regarding each BAC were downloaded from NCBI’s FTP server[9] and curated for quality, focusing on end-sequenced BACs with unique placement of a reasonable length (25 −350 kilobases.) Individual BACs were processed by a Python pipeline written for distributed processing and maps were saved for 233 enzymes, representing all available enzyme recognition sites. The pipeline relies on the Bio.Restriction class from Biopython [12], which identifies restriction digest sites for an enzyme in a sequence. Most of the library is composed of 281,828 end-sequenced BACs from 8 different libraries covering 93% of the genome with an average insert length of 143,311 basepairs. 25,653 insert-sequenced BACs from 37 libraries with an average insert length of 32,309 basepairs are also included. The rare-cutting enzymes were of most interest. To save space, the database was truncated at enzymes which cut a BAC more than 50 times, such digest would create small fragments easily recreated by PCR. Available on the github repository are all the scripts used to generate this library, in case the library should be built locally or updated. Analysis scripts are also available to take advantage of the resource and sequence data related to BACs. These scripts focus on finding maps and sequence for specific BACs and manipulating these maps to design new experiments.

## Acknowledgements

We thank the good people, past and present, associated with and of the Laboratory of Molecular and Computational Genomics, namely Dr. Sam Krerowicz and Dr. Louise Pape. We also thank NHGRI for funding: R21 HG012281.

## References

1. International Human Genome Sequencing C. Initial sequencing and analysis of the human genome. Nature. 2001;409:860. doi:10.1038/35057062 https://www.nature.com/articles/35057062#supplementary-information.

2. H S, B B, UJ K, V M, T S, Y T, et al. Cloning and stable maintenance of 300-kilobase-pair fragments of human DNA in Escherichia coli using an F-factor-based vector. Proceedings of the National Academy of Sciences of the United States of America. 1992;89 18 doi:10.1073/pnas.89.18.8794.

3. Xiao C, Chen Z, Chen W, Padilla C, Colgan M, Wu W, et al. Personalized genome assembly for accurate cancer somatic mutation discovery using tumor-normal paired reference samples. Genome Biology. 2022;23 1:1–34. doi:doi:10.1186/s13059-022-02803-x.

4. Staff N. NCBI retires Clone DB - NCBI Insights. 2019.

5. N O, J B, T E, DB G, BJ K, HH L, et al. Technological challenges and milestones for writing genomes. Science (New York, NY). 2019;366 6463 doi:10.1126/science.aay0339.

6. Kent WJ, Sugnet CW, Furey TS, Roskin KM, Pringle TH, Zahler AM, et al. The Human Genome Browser at UCSC. 2002; doi:10.1101/gr.229102.

7. Eileen T. Dimalanta, Alex Lim, Rod Runnheim, Casey Lamers, Chris Churas, Daniel K. Forrest, et al. A Microfluidic System for Large DNA Molecule Arrays. 2004; doi:10.1021/ac0496401.

8. B T, MS W, S G, K P, S Z, S R, et al. High-resolution human genome structure by single-molecule analysis. Proceedings of the National Academy of Sciences of the United States of America. 2010;107 24 doi:10.1073/pnas.0914638107.

9. EW S, EE B, JR B, K C, J C, DC C, et al. Database resources of the national center for biotechnology information. Nucleic acids research. 2022;50 D1 doi:10.1093/nar/gkab1112.

10. KD B and JT T. Bacterial genomics: the use of DNA microarrays and bacterial artificial chromosomes. Journal of microbiological methods. 2002;49 3 doi:10.1016/s0167-7012(01)00375-x.

11. JC V, JI G, CA H and S V. Synthetic chromosomes, genomes, viruses, and cells. Cell. 2022;185 15 doi:10.1016/j.cell.2022.06.046.

12. Peter J. A. Cock, Tiago Antao, Jeffrey T. Chang, Brad A. Chapman, Cymon J. Cox, Andrew Dalke, et al. Biopython: freely available Python tools for computational molecular biology and bioinformatics. Bioinformatics. 2023;25 11:1422–3. doi:10.1093/bioinformatics/btp163.

